# Interaction of combined stressors and genetic background on recombination rate plasticity in *Drosophila melanogaster*

**DOI:** 10.64898/2026.10.16.759659

**Authors:** Natalia Rivera-Rincón, Emma C. Saurette, Logan G. Havard, Laurie S. Stevison

## Abstract

Recombination is an important source of genetic variation and plays a central role in adaptation and genome stability. Although environmental conditions such as temperature can alter recombination rates, much less is known about how multiple environmental stressors act together. Here, we tested the effects of low temperature and hypoxia, both independently and in combination, on recombination in *Drosophila melanogaster* across five genetic backgrounds. Using a quadruple X-linked marker stock, we measured recombination in 50,209 male backcross progeny from 194 backcrosses exposed to four conditions: control, hypoxia, low temperature, and combined hypoxia and low temperature. Low temperature significantly altered recombination across all three chromosome intervals examined and affected total map length with an average increase of 5 cM. Genetic background strongly influenced the magnitude of altered recombination rate. Hypoxia alone did not significantly change total map length, but its effects differed among genetic backgrounds with significant variation in both the magnitude and direction with an average decrease of 0.96 cM. The response to combined hypoxia and low temperature was not simply the sum of the responses to each stressor alone, with an average increase of 6.78 cM, that also varied across genetic backgrounds in magnitude and direction as well as over time. DGRP_57 and DGRP_508 showed some of the largest increases in recombination under combined stress, whereas DGRP_491 showed a decrease. DGRP_391 was robust across all treatments. Despite these changes in crossover frequency, crossover interference remained largely stable across treatments, with only the *y-cv-v* interval pair showing a significant treatment-by-genotype interaction. Together, these results show that recombination responses to environmental stress depend strongly on genetic background and on the combination of stressors experienced. Our findings highlight the importance of considering both genetic variation and interacting environmental conditions when predicting how populations may respond to increasingly complex environmental change.

## Introduction

Recombination is a crucial process that occurs during meiosis, where homologous chromosomes exchange genetic material, leading to the formation of genetically diverse gametes (Stapley et al. 2017; Stevison et al. 2017). This process is important because it ensures that offspring inherit a unique combination of genes from both parents, contributing to genetic diversity within a population. Genetic diversity is key to the ability of species to adapt to changing environments, as it increases the likelihood that some individuals will possess traits suited to new or altered conditions (Nachman 2002; Lloyd et al. 2018; Samuk et al. 2020; Rybnikov et al. 2023). Recombination not only facilitates the shuffling of alleles but also enables the repair of damaged DNA and the removal of harmful mutations (Charlesworth 1993). Given its central role in inheritance and evolution, understanding how recombination rates can vary in response to environmental factors is of great significance (Zhong and Priest 2010).

The phenomenon of recombination rate plasticity, in which recombination rates change in response to environmental conditions, has been a subject of interest in Drosophila research for over a century. Early studies, such as those by Plough (1917, 1921), demonstrated that heat stress significantly alters recombination rates during meiosis, providing some of the first evidence of environmental influence on genetic mechanisms. These observations were extended by Graubard (1932), who examined the effects of temperature in the context of chromosomal inversions and showed that the recombination response to temperature can depend on chromosome structure, with inversion heterozygosity strongly suppressing crossing over despite changes in temperature. Graubard (1934) subsequently examined temperature effects on crossing over and interference more directly, further supporting the sensitivity of meiotic recombination to thermal conditions. Subsequent research has shown that recombination rate plasticity is not limited to temperature but can also occur in response to other environmental factors. For example, nutritional stress, as explored by Neel (1941), was found to influence recombination rates, suggesting that energy availability plays a critical role in regulating this process. Similarly, mating conditions, as investigated by Zhong and Priest (2010), were shown to modulate recombination rates, further illustrating the range of environmental and reproductive factors capable of altering recombination.

In addition to thermal and nutritional stress, hypoxia has emerged as another environmental factor associated with variation in recombination. Aggarwal et al. (2015) showed that long-term selection for hypoxia tolerance altered recombination rates in Drosophila, suggesting that recombination can evolve in response to sustained low oxygen conditions. These findings align with broader observations of hypoxia’s effects on cellular and physiological processes, such as metabolism (Zhou et al. 2008; Van Voorhies 2009) and gene expression (Azad et al. 2009, 2012). Hypoxia can also generate reactive oxygen species and oxidative stress (Zhou et al. 2007). Hypoxia and oxidative stress show substantial overlap in their transcriptional responses and are both hallmarks of several cancers (Suzuki et al. 2021; Bae et al. 2024), which collectively show the profound influence of oxygen availability on organismal biology. In *D. melanogaster*, studies have identified genetic variation in tolerance to oxidative damage in the Drosophila Genetic Reference Panel (DGRP) (Mackay et al. 2012; Weber et al. 2012), paralleling genotypic variation in hypoxia response (Mossman et al. 2017). However, oxidative stress does not necessarily translate into increased recombination; paraquat-induced oxidative damage in honeybees produced no overall increase in meiotic recombination rate (Langberg et al. 2018).

Despite these advances, the combined effects of oxygen availability and cold stress on recombination rates remain poorly understood. The interaction between these stressors is particularly relevant in the context of climate change, where exposure to low temperatures may coincide with hypoxic conditions in certain habitats, such as overwintering sites or microenvironments with limited oxygen availability.

Studies on the synergistic or antagonistic effects of combined stressors have provided insights into other biological processes, such as metabolic rate suppression under hypoxia and cold (Boardman et al. 2016), yet the implications for recombination rate plasticity remain unexplored.

Together, these studies highlight the importance of genetic background in determining responses to environmental stressors. Individuals with different genetic backgrounds can differ in their baseline recombination rates, their sensitivity to stressors, and the molecular pathways they utilize to cope with environmental challenges (Comeron et al. 2012). For example, differences in stress-response genes, metabolic pathways, or chromosomal architecture across genetic backgrounds could lead to variability in how recombination rates are modulated (Hoffmann and Parsons 1991). Including different genetic backgrounds, provides a more comprehensive understanding of how genetic and environmental factors interact to shape recombination plasticity.

Drosophila is an ideal model organism for this kind of study, and it is particularly advantageous due to its well-characterized genetic system, rapid life cycle, and the availability of numerous tools for manipulating genetic and environmental variables. Previous studies have provided knowledge on the effects of stressors such as acute heat exposure (Jackson et al. 2015), cold stress (Zhong and Priest 2010), and hypoxia (Aggarwal et al. 2015) on Drosophila recombination rate. Building on this foundation, here we aimed to investigate the combined effects of hypoxia and cold stress on recombination rates, filling a critical gap in our understanding of how organisms might respond to the multifaceted challenges posed by climate change. To address this gap, and as part of a companion study characterizing the physiological and reproductive effects of these same stressors (Rivera-Rincón et al. 2026), classic genetic crosses were conducted using a quadruple X-linked chromosome mutant stock and five different genetic backgrounds. These experimental crosses allowed for the calculation of recombination rates across three distinct genetic intervals, for five different genetic backgrounds. Our work provides growing evidence that plasticity in recombination is highly variable across genetic backgrounds.

## Methods

The crosses and genetic backgrounds used here were previously described in a companion study of the physiological and reproductive effects of hypoxia and low temperature (Rivera-Rincón et al. 2026). The fly husbandry, treatment, and crossing methods are described below and are summarized in Figures 1-2. Male offspring from the previously described backcrosses were analyzed here for evidence for variation in meiotic crossing over.

**Figure 1.**
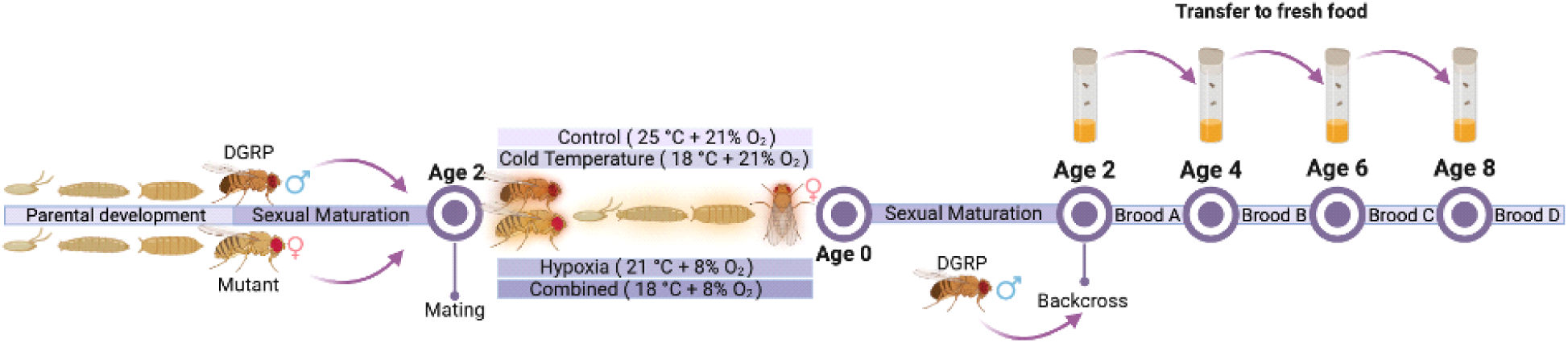
Experimental Design. Graphical representation of the three-generation cross design with the ages at which crosses and multiday transfers were made. Age is defined as days post-eclosion. Drosophila Genome Reference Panel (DGRP) indicates one of the five strains used across these four perturbations. A companion study measured reproduction and organismal physiology from the same experiment (Rivera-Rincón et al. 2026). Here, we report changes in meiotic recombination rate using a mutant strain with four visible markers.

**Figure 2.**
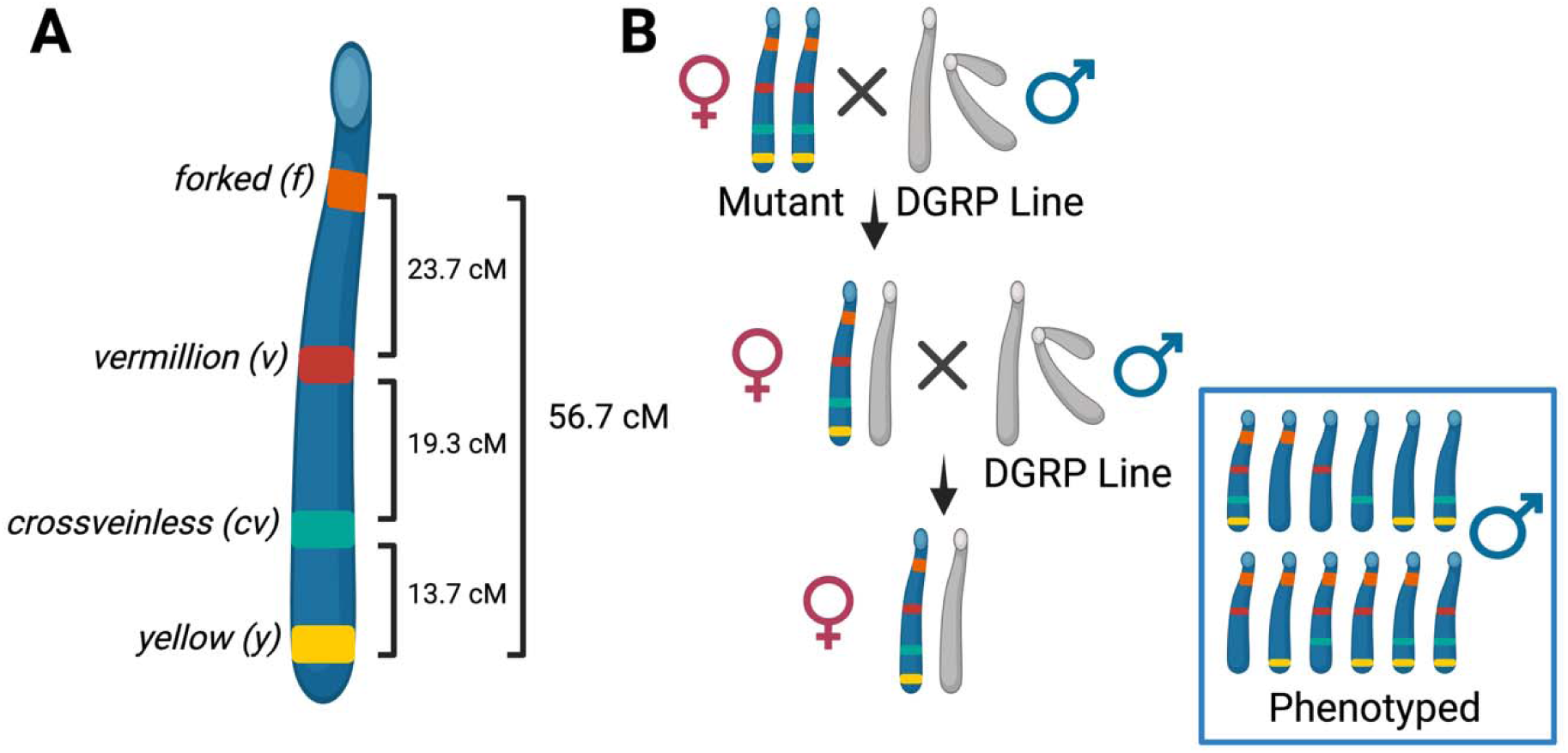
Graphical description of the visible markers used, and the genetic distances between the intervals evaluated (A), and a cartoon schematic of the genetic crosses and backcrosses design (B). Only male progeny with visibly expressed mutant phenotypes were assayed for measures of meiotic recombination rate.

### Stocks, Fly husbandry, and Genetic Crosses

We used a mutant stock with four X-linked visible markers to measure changes in recombination rate. Specifically the markers *yellow (y), cross-veinless (cv), vermillion (v),* and *forked (f)* were used to compare recombination frequency (Figure 2A), which have been previously used to examine recombination rate plasticity (Novak et al. 2026). We also used five Drosophila Genetic Reference Panel (DGRP) stocks: DGRP_42 (BDSC_25193), DGRP_57 (BDSC_29652), DGRP_391 (BDSC_25191), DGRP_491 (BDSC_28202), and DGRP_508 (BDSC_28205) --obtained from the Bloomington Drosophila Stock Center (Research Resource Identifiers in parentheses; NIH P40OD018537). These stocks were selected based on previously reported paraquat-survival and chill-coma-recovery assays conducted with the DGRP panel (Mackay et al. 2012; Weber et al. 2012), as described in the companion study of the physiological and reproductive effects of these same stressors (Rivera-Rincón et al. 2026, see Figure S1). Specifically, these five stocks cover the range of values across both distributions of responses.

All stocks were maintained under standard DGRP-panel husbandry conditions (25°C, 12:12 light:dark photoperiod in a Percival DR41-VL incubator, standard cornmeal-sugar-yeast-agar media in polypropylene enclosures). This was also the control condition throughout the experiment and whenever a treatment was not being directly applied as shown in Figure 1.

A range of 10-24 replicate crosses were set up per genetic background per treatment combination, for a total of 281 crosses. An average of 10 quadruple X-linked mutant females and 5 age-matched DGRP males (2 days old, post eclosion) were crossed (Figure 2B). These F_1_ crosses were placed on one of the four different treatment combinations: control, hypoxia, low temperature, or combined (Figure 1). The F_1_ females were collected within 8 hours of eclosing and held virgin for two to three days for sexual maturation before being backcrossed to age-matched DGRP males. The DGRP males were backcrossed to the F_1_ to have a built-in fail-safe for the X-link mutant crosses (Novak et al. 2026). This resulted in 195 backcrosses across genetic background and treatments, of which 194 retained at least five scored male progeny and were used for recombination-rate models (see Data Quality Control). Each backcross was transferred to fresh food every 48-hours to prevent overcrowding of the vials, and to partition the recombination rates based on age of the female parent. This separated each cross into four different timepoints (or broods): A (Day 2-3), B (Day 4-5), C (Day 6-7), and D (Day 8-9) (Figure 1). Only male offspring from the backcrosses were phenotyped (Figure 2B), and this was done over a seven-day period starting when flies began eclosing from each vial.

### Treatments

Low temperature and oxygen levels were previously described in the companion study of these same treatments (Rivera-Rincón et al. 2026, see Figure S2). Briefly, the control temperature was 25□, and the stressful low temperature was chosen based on a pilot experiment to be 18□. Oxygen levels of 21% O_2_ (normoxia) and 8% O_2_ (hypoxia) were chosen based on prior work showing comparable reductions in survivorship between the two stressors (Zhou et al. 2007). While previous hypoxia studies in Drosophila have generally exposed flies only briefly, intermittently, or during a single developmental stage (Zhou et al. 2008, 2011; Azad et al. 2009; Weber et al. 2012; Zhou and Haddad 2013; Callier et al. 2015; Wang et al. 2017; Stobdan et al. 2024), here, we instead exposed flies to hypoxia throughout the entire developmental cycle (egg-to-adult) to study the impact on meiotic recombination. Oxygen concentration was maintained using BioSpherix chambers housed in temperature-controlled incubators; airflow (21% or 8% certified O_2_, AirGas) was flushed and refreshed for 20 minutes every 12 hours and after each chamber opening. A small USB-powered fan was also added to each chamber to circulate air throughout the chambers between flushing.

### Data Quality Control

Prior to the recombination rate analysis, we assessed the raw phenotype data for potential differences in scoring among observers, consistent with previous studies where multiple observers were involved in data collection (Schüßler et al. 2024; Novak et al. 2026). Here, eight observers scored crossover phenotypes during the experiment. Because triple crossovers (TCOs) are expected to occur at low frequencies, unusually high TCO rates were used to identify possible scoring inconsistencies. For each observer, we compared their TCO rate with the distribution across all eight observers and flagged rates that were more than one median absolute deviation (MAD) above the overall mean.

This identified one observer with a TCO rate of 9.3%, which was 15.4 times higher than expected value of 0.605% based on FlyBase map distances (Comeron et al. 2012), and more than three times higher than the mean TCO rate of the remaining observers. A second observer was also excluded because they scored only eight males in total, providing too little data for reliable comparison.

As done in another recent study (Novak et al. 2026), we also tested whether the number of males scored per vial differed in variability among observers. In the raw dataset, a Levene’s test showed significant heterogeneity of variance among observer × backcross groups (F_863,4667_ = 1.35, p = 9.5 × 10□¹□). After removing the two flagged observers, this heterogeneity was no longer detected (F_668,2228_ = 1.01, p = 0.45), indicating that the remaining observations were more consistent across scorers.

Removing these two observers excluded 2,634 of 5,531 vial-day records, corresponding to 35,304 of the 75,434 males originally scored. We also excluded nine crosses showing evidence of contamination, either because larvae or pupae were present in vials expected to contain virgins or because mutant female offspring appeared in the backcross progeny. The final dataset contained 50,209 male backcross progeny from 195 backcrosses. Of these, 194 backcrosses retained at least five scored males per vial and were included in the recombination-rate analyses (Table 1).

**Table 1.** Sample Sizes across experiment. Total numbers of F1 crosses and backcrosses set up for each genotype and treatment combination. The respective total number of offspring and number of male progeny per treatment, per genetic background is included as well as the totals at the bottom. Only males were used for recombination rate analysis.

| **Genetic Background** | **Treatment** | **F<sub>1</sub> replicate crosses** | **Backcrosses** | **Total Offspring** | **Male Progeny** |
| --- | --- | --- | --- | --- | --- |
| **42** | Control | 11 | 7 | 3302 | 1625 |
|  | Hypoxia | 16 | 10 | 5032 | 2550 |
|  | LowTemp | 14 | 11 | 4348 | 2105 |
|  | Combined | 17 | 12 | 6040 | 3087 |
| **57** | Control | 13 | 10 | 6117 | 2521 |
|  | Hypoxia | 12 | 8 | 4273 | 1790 |
|  | LowTemp | 16 | 11 | 8504 | 3906 |
|  | Combined | 11 | 9 | 5117 | 2289 |
| **391** | Control | 24 | 10 | 10190 | 4074 |
|  | Hypoxia | 12 | 8 | 4025 | 1508 |
|  | LowTemp | 15 | 11 | 5173 | 2077 |
|  | Combined | 19 | 12 | 5821 | 2265 |
| **491** | Control | 12 | 10 | 4964 | 2107 |
|  | Hypoxia | 10 | 6 | 3649 | 1527 |
|  | LowTemp | 17 | 12 | 7503 | 3475 |
|  | Combined | 12 | 8 | 5760 | 2455 |
| **508** | Control | 15 | 10 | 7159 | 2904 |
|  | Hypoxia | 11 | 8 | 5251 | 2225 |
|  | LowTemp | 11 | 11 | 5741 | 2577 |
|  | Combined | 13 | 11 | 7075 | 3142 |
| **TOTALS** |  | 281 | 195 | 115044 | 50209 |

### Inference of Meiotic Recombination Rate

Recombination rates were analyzed using a classic backcross approach with four previously mapped X-linked visible genetic markers: *yellow* (*y*) which is a mutation that leads to lighter yellow bodied flies, *crossveinless* (*cv*), which has loss of the crossveins on the wings, *vermilion* (*v*), which has a bright red eye, and *forked* (*f*), which is a mutation of the bristles (Figure 2A), as described by Lindsley and Grell (1967). These markers allowed for the identification of crossover events between three specific genomic intervals (*y–cv, cv–v*, and *v–f*), and the estimation of recombination frequencies which was converted to total map length via a Kosambi mapping function (Kosambi 1944). Recombination events were categorized in four different types based on the number of recombination events: non-crossover (NCO), single crossover (SCO, y–cv, cv–v, or v–f), double crossover (DCO), and triple crossover (TCO). For each vial, the number of crossover events was determined by summing the observed single, double, and triple crossovers. Interval-specific recombination rates were calculated separately for *y–cv, cv–v*, and *v–f* by dividing the number of recombinants in each interval by the total number of males. The total map length was estimated as the sum of recombination rates across all three intervals.

### Crossover Interference (COI)

Coefficient of coincidence (COC) and crossover interference were calculated for the two pairs of adjacent intervals (*y-cv* and *cv-v*; *cv-v* and *v-f*) and across all three intervals (*y-cv-v-f*). For each pair of adjacent intervals, we first calculated the number of double crossovers (DCOs) expected if crossovers in the two intervals occurred independently. This was done by multiplying the crossover frequencies of the two intervals by the total number of male progeny scored for each vial-by-brood combination. The same approach was used to calculate the expected number of triple crossovers (TCOs) across all three intervals.

The coefficient of coincidence was calculated by dividing the observed number of DCOs or TCOs by the corresponding expected number. Crossover interference was then calculated as 1 − COC, with larger values indicating stronger interference.

### Statistical analysis

Binomial generalized linear mixed-effects models (GLMMs) were used to analyze recombination separately for each of the three chromosome intervals and across the total map length. For each analysis, the response was the proportion of offspring showing a crossover event. Treatment (control, hypoxia, low temperature, or combined stress), genetic background, and brood were included as fixed effects, along with their interactions. Brood represented time since mating and was divided into four two-day intervals. Backcross vial was included as a random effect to account for repeated observations from the same cross.

Models were fit in R using glmer from the ‘lme4’ package, with crossover and non-crossover counts specified as a binomial response. The bobyqa/nlminb optimizer was implemented through the ‘optimx’ package when needed to improve model convergence.

For total map length, we also separated Treatment into its oxygen and temperature components to test the individual effects of hypoxia and low temperature, as well as whether their effects depended on one another (Supplementary Table S1).

Interference was analyzed separately for each of the three interval combinations using generalized linear models with a quasi-Poisson error distribution and log link. The observed number of DCOs or TCOs was used as the response variable, with Treatment, Genetic Background, and their interaction included as fixed effects. The log of the expected DCO or TCO count was included as an offset, facilitating a direct test of whether the number of observed crossovers differed from what would be expected based on the crossover frequencies of the individual intervals. Due to small sample size within each timepoint, timepoint was not considered in the statistical analysis. Timepoints with small counts such that the expected number of TCOs were zero were removed.

When significant effects were detected, estimated marginal means (EMMs) were used for post hoc comparisons among treatments within each genetic background and brood. Statistical analyses were performed in R v4.5.3 (R Core Team 2025) using ‘lme4’ (v.1.1-34, Bates et al. 2015), ‘emmeans’ (v.1.10.2, Lenth and Piaskowski 2025), and ‘car’ (v.3.1-2, Fox et al. 2012). Figures 1-2 were created using BioRender and Figure 3-4 as other supplemental plots were generated with ‘ggplot2’ (Wickham 2011).

**Figure 3.**
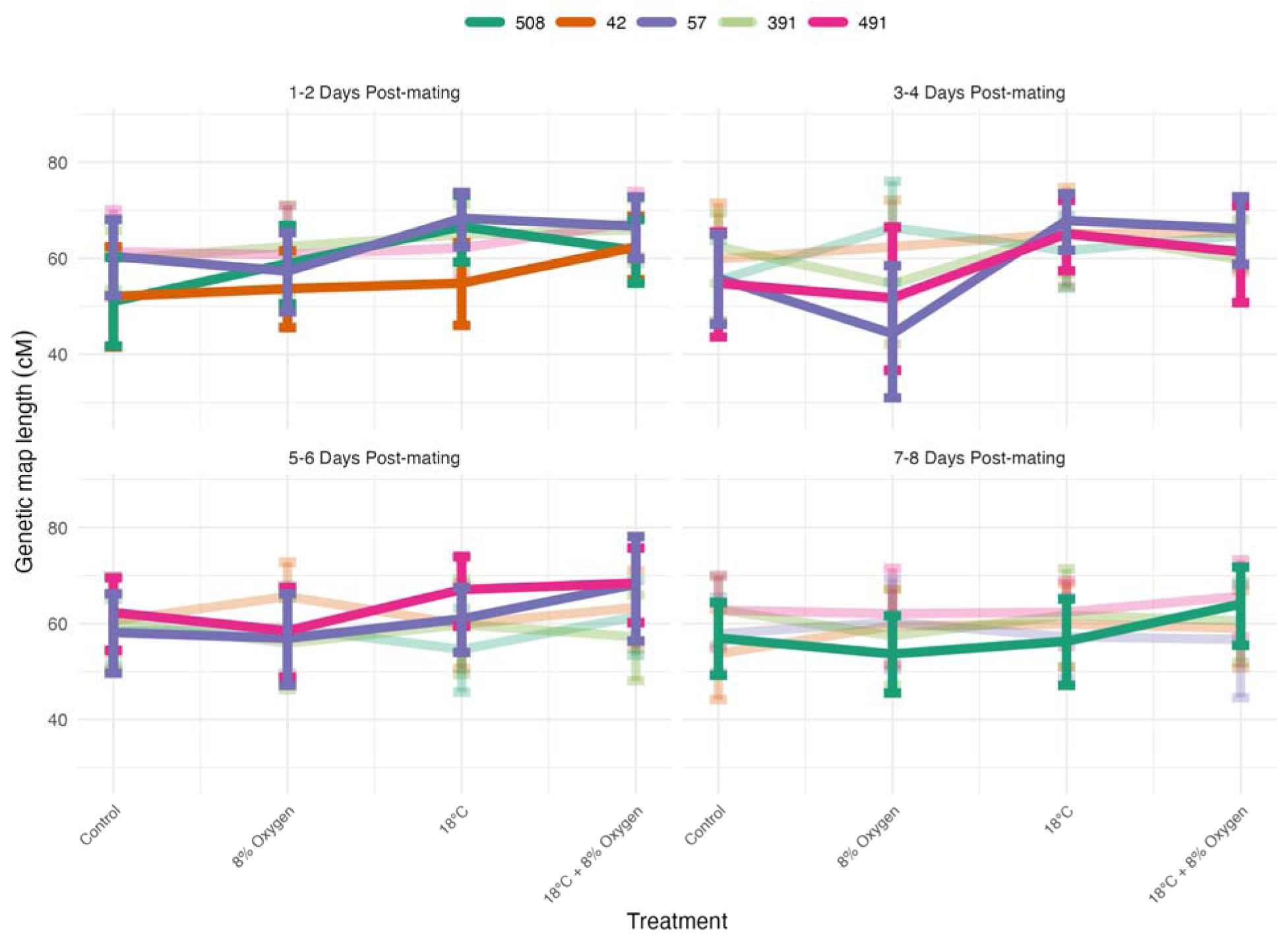
Genetic map length estimates vary across genotypes and treatments and over time. To highlight the 3-way interaction for strain, treatment, and brood, the emmeans were extracted from the full model and were plotted for total genetic map length across the experiment. Model table results shown in *Table 3*. Significance of each genetic background indicated by intensity of the line color. Full posthoc results per strain by time point for each treatment contrast can be found in Supplemental Figures 3-7. A stacked bar plot showing these results by interval can be found in Supplementary Figure 2 and further split by timepoint in Supplementary Figure 8.

**Figure 4.**
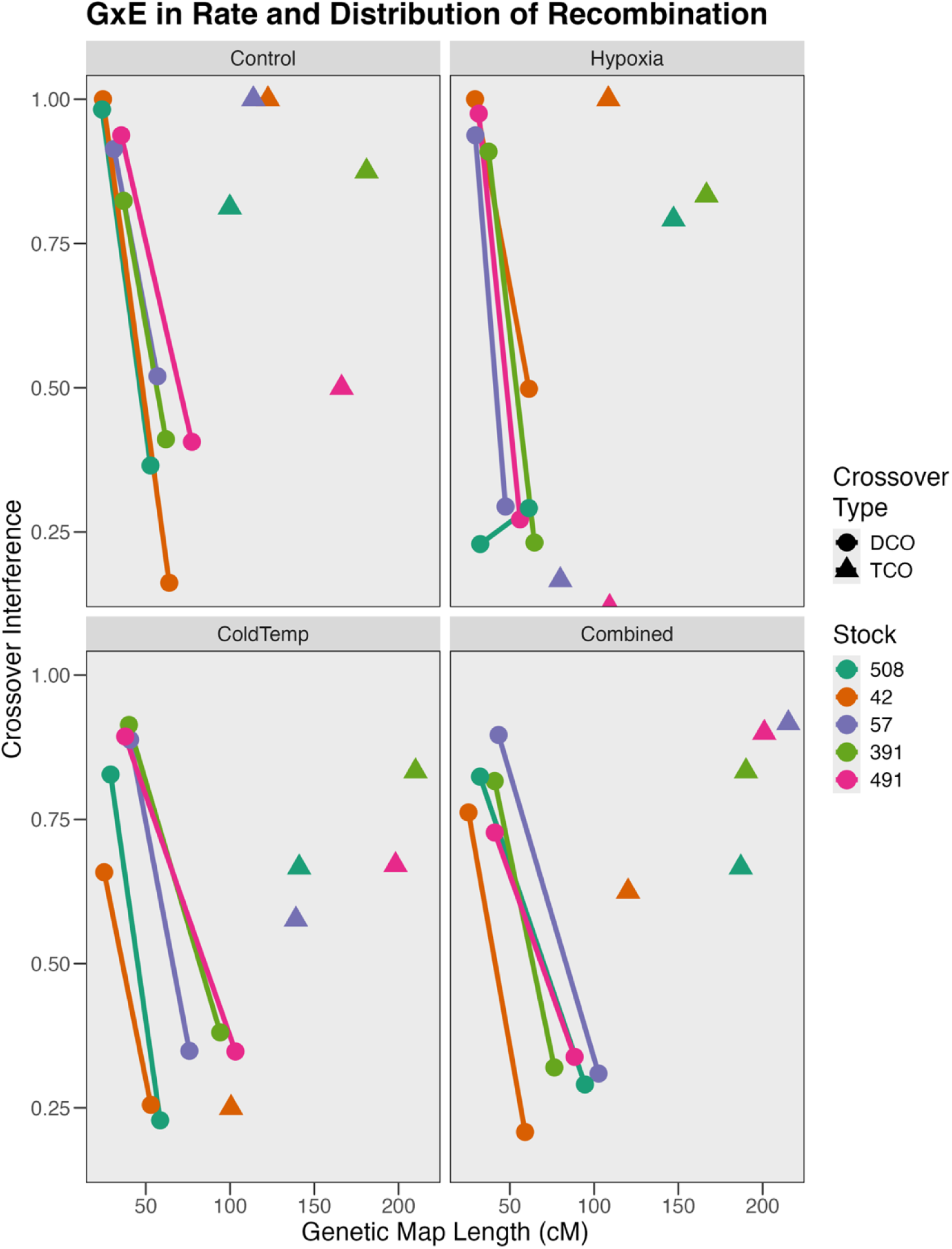
Genetic Distance versus Interference. The relationship between recombination rate and interference follows an expected decay function with interference being higher in intervals nearest to the telomere. For the control treatment (top left), the five genotypes cluster along both axes. The hypoxia treatment (top right) shows decreased recombination, as compared to coldtemp, where the genotypes are more spread along their lengths (bottom left). Finally, the combined treatment shows an intermediate change between the two treatments independently. Crossover interference was examined separately based on double crossovers (DCOs) from pairs of intervals, which are connected by a line, and triple crossovers (TCOs) from all three intervals combined.

## Data Availability

A snapshot of the code at the time of publication can be found in the following GitHub repository https://github.com/StevisonLab/Recombination_Hypoxia_LowTemp_in_DMEL (https://doi.org/10.5281/zenodo.23401297). This experiment was done in conjunction with an analysis of reproduction and physiological effects of these same treatments and genotypes. The corresponding data and code can be found in the associated GitHub repository https://github.com/StevisonLab/Combination-effects-of-hypoxia-and-low-temperature-in-Drosophila/.

## Results

After quality control, the final dataset included 115,044 offspring from 281 crosses and 194 backcrosses spanning five genetic backgrounds and four treatment conditions. Recombination rates were calculated using phenotypes from 50,209 male offspring. Of the 195 backcrosses, 194 retained at least five scored males and were included in the recombination-rate models (Table 1). Recombination was analyzed separately for each of the three chromosome intervals and across the total map length (Table 2). Additionally, crossover interference was measured using adjacent intervals and comparison of the observed double and triple crossover events to the expected number of events (Table 3).

**Table 2.**
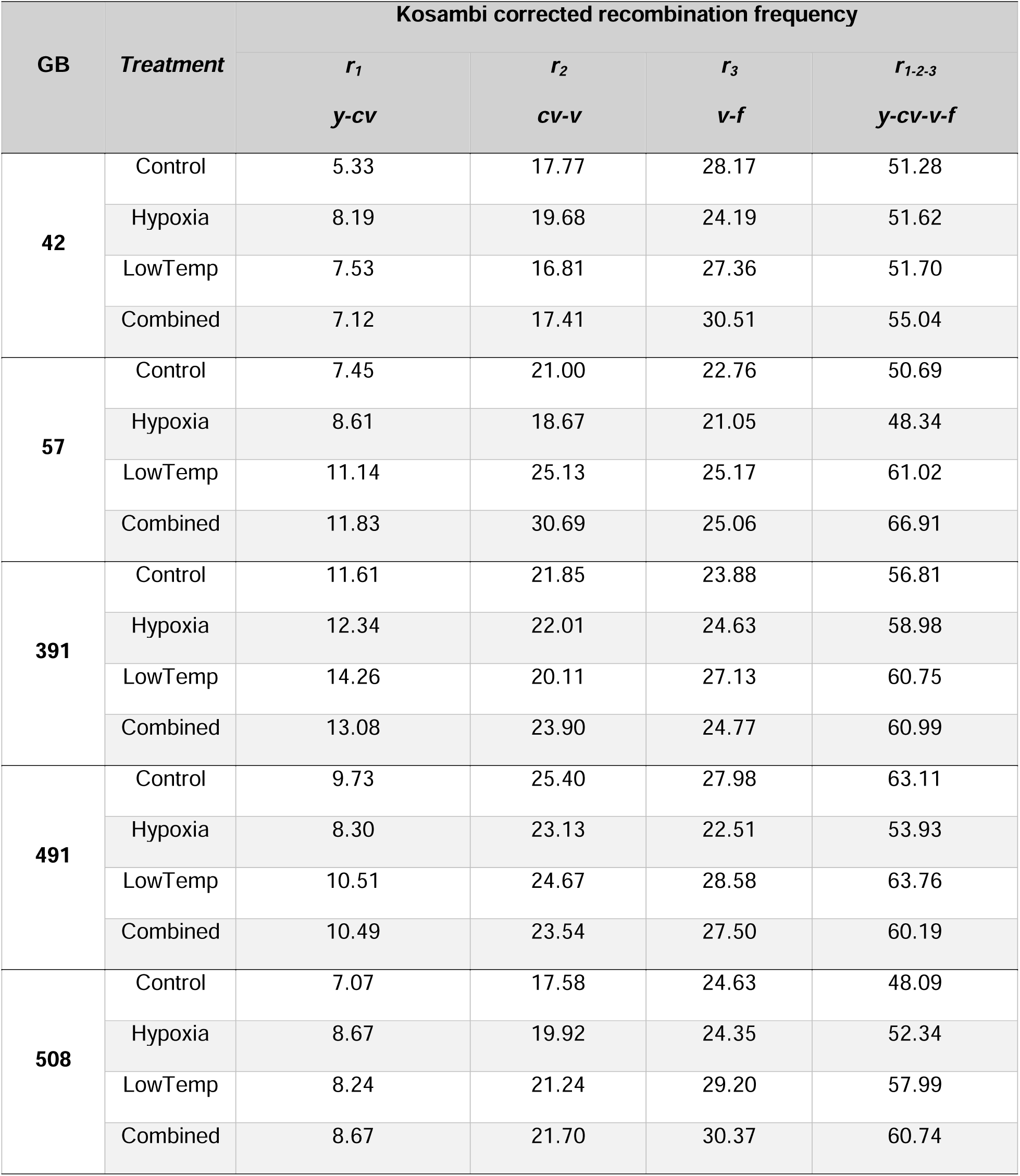
Recombination rate results. Mean Kosambi corrected recombination frequencies were summarized across each marker interval, genetic background, and treatment and across the four time points (see *Figure 1*).

**Table 3.** Crossover Interference Results. Expected double crossovers were estimated based on the product of adjacent intervals with the coefficient of coincidence (c.o.c.) calculated from the ratio of the observed double crossovers to expected double crossovers between adjacent intervals. Interference was estimated as 1-c.o.c. Shown below are mean values across time points.

| Crossover Interference (1-coefficient of coincidence) |  |  |  |  |
| --- | --- | --- | --- | --- |
| GB | Treatment | c<sub>12</sub> | c<sub>23</sub> | c<sub>13</sub> |
|  |  | y-cv-v | cv-v-f | y-cv-v-f |
| 42 | Control | 1.00 | 0.16 | 0.25 |
|  | Hypoxia | 1.00 | 0.50 | 0.63 |
|  | LowTemp | 0.66 | 0.26 | 1.00 |
|  | Combined | 0.76 | 0.21 | 1.00 |
| 57 | Control | 0.91 | 0.52 | 0.58 |
|  | Hypoxia | 0.94 | 0.29 | 0.92 |
|  | LowTemp | 0.89 | 0.35 | 1.00 |
|  | Combined | 0.90 | 0.31 | 0.17 |
| 391 | Control | 0.82 | 0.41 | 0.83 |
|  | Hypoxia | 0.91 | 0.23 | 0.83 |
|  | LowTemp | 0.91 | 0.38 | 0.88 |
|  | Combined | 0.82 | 0.32 | 0.83 |
| 491 | Control | 0.94 | 0.41 | 0.67 |
|  | Hypoxia | 0.98 | 0.27 | 0.90 |
|  | LowTemp | 0.89 | 0.35 | 0.50 |
|  | Combined | 0.73 | 0.34 | 1.00 |
| 508 | Control | 0.98 | 0.37 | 0.67 |
|  | Hypoxia | 0.23 | 0.29 | 0.67 |
|  | LowTemp | 0.83 | 0.23 | 0.81 |
|  | Combined | 0.82 | 0.29 | 0.79 |

Treatment significantly affected recombination in all three intervals and across the total map length (Table 4). Genetic background also had a significant effect in every interval and for total map length. Brood, which reflects time since mating, significantly affected recombination in the *y-cv* and *v-f* intervals. In addition, the effects of treatment changed across broods and genetic backgrounds. The three-way Treatment × Genetic Background × Brood interaction was significant for all three intervals and total map length (Table 4), showing that the response to treatment depended on both genetic background and time since mating (Figure 3). Crossover interference was largely consistent across genotypes and intervals, with higher interference in intervals spanning chromosome ends as compared to more central intervals (Table 4). Comparison of changes in both crossover frequency and distribution across treatments and genotypes visually reveals the variation in these changes (Figure 4).

**Table 4.** Summarized model table for each interval and for total map length for recombination rate (r) and crossover interference (c). Values represent p-values for each model parameter with bolded values representing statistical significance. Due to the small number of double crossovers, interference was calculated as the sum across all replicates of each time point. Therefore, ‘Brood’ was not included in the model for Interference. Full model tables can be found in the GitHub Repository for each interval as well as post hoc tables.

| Model Parameter | Df | r<sub>1</sub> | r<sub>2</sub> | r<sub>3</sub> | r<sub>1-2-3</sub> | c<sub>12</sub> | c<sub>23</sub> | c<sub>13</sub> |
| --- | --- | --- | --- | --- | --- | --- | --- | --- |
|  |  | y-cv | cv-v | v-f | y-cv-v-f | y-cv-v | cv-v-f | y-cv-v-f |
| Treatment | 3 | **2.91E-8** | **8.76E-5** | **9.30E-8** | **1.49E-8** | 0.16 | 0.17 | 0.36 |
| Genetic Background<br>(GB) | 4 | **2.43E-8** | **2.14E-8** | **2.10E-3** | **1.30E-2** | 0.21 | 0.18 | 0.93 |
| Brood | 3 | **4.45E-5** | 0.223 | **9.85E-3** | 0.112 | Na | Na | Na |
| Treatment:GB | 12 | 0.052 | **0.021** | 0.404 | 0.123 | **6.23E-3** | 0.21 | 0.23 |
| Treatment:Brood | 9 | **7.64E-7** | **0.019** | **7.37E-3** | **1.97E-4** | Na | Na | Na |
| GB:Brood | 12 | **3.38E-8** | **1.94E-3** | **9.09E-7** | **6.58E-8** | Na | Na | Na |
| Treatment:GB:Brood | 36 | **2.28E-4** | **6.69E-3** | **0.024** | **6.55E-5** | Na | Na | Na |

When Treatment was separated into its oxygen and temperature components for the total map length analysis, temperature had a strong effect on recombination (χ²□ = 39.36, p = 3.53 × 10□¹□), whereas oxygen alone did not (χ²□ = 0.85, p = 0.36). However, the effect of oxygen differed among genetic backgrounds (Oxygen × Genetic Background: χ²□ = 9.56, p = 0.048). We also detected a significant interaction among oxygen, temperature, genetic background, and brood (χ²□□ = 27.27, p = 0.0071; Supplementary Table S1), further showing that the response to the two stressors depended on both genotype and time since mating.

The Oxygen × Temperature interaction itself was not significant for total map length (χ²□ = 0.94, p = 0.33). This suggests that there was no single, consistent combined effect of hypoxia and low temperature across all flies. Instead, the way the two stressors acted together varied among genetic backgrounds and across broods (Supplementary Table S1).

### Genetic background influences differences in responses to combined stressors

Responses to treatments differed substantially among genetic backgrounds (Table 2; Figure 3). Some backgrounds showed clear increases in recombination, while others changed only slightly, and the direction and magnitude of the response often differed across chromosome intervals.

DGRP_57 showed the strongest response of the five genetic backgrounds (Figure S4). Under the combined hypoxia and low-temperature treatment, recombination increased in all three intervals relative to the control (Figure 3). The *y-cv* interval increased from 7.45 to 11.83 cM (+4.38 cM), while the *cv-v* interval showed the largest increase, from 21.00 to 30.69 cM (+9.69 cM; Figure S2). A smaller increase was also observed in the *v-f* interval, from 22.76 to 25.06 cM (+2.30 cM). Together, these changes produced an increase in total map length from 50.69 to 66.91 cM (+16.21 cM), the largest increase observed among the five backgrounds (Table 2). In contrast, hypoxia alone slightly reduced total map length in DGRP_57 by 2.35 cM (Table 2).

DGRP_42 showed a much weaker response to the treatments (Figure S3). Recombination in the *y-cv* interval increased modestly under all three stress treatments relative to the control (+1.8 to +2.9 cM), whereas changes in the cv-v interval were small and varied in direction, ranging from a 0.96 cM decrease under low temperature to a 1.91 cM increase under hypoxia (Table 2). The *v-f* interval changed little except under the combined treatment, where it increased by 2.34 cM (Figure S2). Overall, these interval-specific changes had little effect on total map length, which increased only from 51.28 cM in the control to 55.04 cM under combined stress (+3.77 cM; Table 2). Importantly, the total map length for DGRP_42 in the control (51.28 cM) is extremely close to the reported control map length for this same strain in a recent study from our lab using a similar crossing scheme, but investigating dietary perturbation (51.93 cM; Novak et al. 2026), suggesting reproducibility across studies. Interestingly, recombination rate in DGRP_42 was much higher under caloric restriction (Novak et al. 2026), despite its robustness to the treatments used in this study, suggesting the plasticity varies significantly across perturbations.

DGRP_508 showed a more consistent increase in recombination. All three intervals increased under each stress treatment relative to the control, with the strongest response again occurring under combined hypoxia and low temperature. Under this treatment, the *y-cv*, *cv-v*, and *v-f* intervals increased by 1.60, 4.12, and 5.74 cM, respectively (Figure S2). Total map length increased from 48.09 to 60.74 cM (+12.65 cM), the second-largest increase among the five genetic backgrounds (Table 2).

DGRP_491 was the only genetic background in which most treatments reduced total map length relative to the control (Figure S6). Hypoxia produced the strongest decrease, reducing map length by 9.18 cM, the largest decrease observed for any single treatment in the study. The combined treatment also reduced map length by 2.92 cM, while low temperature alone produced only a small increase of 0.65 cM. DGRP_391 showed a different pattern, with modest increases across all three treatments (Figure S5). Its largest change occurred under the combined treatment, where total map length increased from 56.81 to 60.99 cM (+4.18 cM; Table 2).

Overall, the response to stress differed considerably among genetic backgrounds. Combined hypoxia and low temperature produced the largest increases in DGRP_57 and DGRP_508, more moderate increases in DGRP_391 and DGRP_42, and little or negative change in DGRP_491. These differences show that genetic background plays an important role in determining both the magnitude and direction of recombination responses to stress (Figure 4).

### Low temperature response enhanced when combined with hypoxia

Comparing the independent low temperature treatment to the combined hypoxia and low-temperature treatment revealed clear differences among genetic backgrounds and chromosome intervals. In DGRP_57, low temperature alone increased total map length by 10.33 cM relative to the control, from 50.69 to 61.02 cM (Table 2). Much of this increase came from the *cv-v* (+4.14 cM) and *y-cv* (+3.69 cM) intervals (Figure S2). Under the combined treatment, total map length increased even further, reaching 66.91 cM. The strongest change occurred in the *cv-v* interval, which increased by 9.69 cM relative to the control, more than twice the increase observed under low temperature alone. Although the overall Treatment × Genetic Background interaction was not significant (Table 4), this pattern suggests that adding hypoxia strongly exaggerated the response to low temperature in DGRP_57.

A similar, although less pronounced, pattern was seen in DGRP_508. In the *v-f* interval, recombination increased by 4.57 cM under low temperature alone and by 5.74 cM under the combined treatment (Figure S2). This contributed to a total map length of 60.74 cM under combined stress, an increase of 12.65 cM relative to the control and the largest value observed for this genetic background.

Together, these patterns suggest that low temperature can alter recombination on its own, but the addition of hypoxia can strengthen that response in some genetic backgrounds. The effect was not uniform across genotypes or chromosome intervals, reinforcing the importance of genetic background in shaping recombination responses to multiple environmental stressors.

Overall, the response to stress differed considerably among genetic backgrounds. Combined hypoxia and low temperature produced the largest increases in DGRP_57 and DGRP_508, more moderate increases in DGRP_391 and DGRP_42, and little or negative change in DGRP_491. These differences show that genetic background plays an important role in determining both the magnitude and direction of recombination responses to stress (Figure 4).

### Crossover Interference Is Largely Unaffected by Treatment

Although the treatments produced clear changes in recombination rate and genetic map length, crossover interference remained relatively stable (Table 3). Of the three interval combinations tested, only the *y-cv* and *cv-v* pair showed a significant Treatment × Genetic Background interaction (quasi-Poisson GLM, F□□,□□ = 2.66, p = 0.0062). No significant effects of Treatment, Genetic Background, or their interaction were detected for the *cv-v* and *v-f* interval pair (F□□,□□ = 1.37, p = 0.21) or for triple crossovers across the full *y-cv-v-f* region (F□□,□□ = 1.33, p = 0.23, Figure 4).

For the *y-cv* and *cv-v* interval pair, a pairwise post hoc comparison between treatment groups identified only one significant difference for DGRP_508, which showed an increase in interference between the hypoxia and the combined treatment groups, (log-scale contrast estimate = −1.86, p = 0.025, Figure 4).

Across genetic backgrounds and treatments, interference generally decreased as genetic map length increased, but this relationship did not show a consistent shift among treatments. Overall, these results suggest that hypoxia and low temperature can change how often and where crossovers occur without strongly altering crossover interference.

## Discussion

The variation among genetic backgrounds observed in this study indicates that recombination rate does not respond to environmental stress in a predictable direction. In some cases, even neighboring chromosome intervals differed in the direction or magnitude of their response, suggesting that broader genetic intervals may mask finer-scale changes in recombination. More detailed mapping approaches could help identify where these differences arise and whether particular genomic regions are especially sensitive to environmental conditions (Stevison et al. 2017). Together, these results add to growing evidence that recombination is a flexible process shaped by both environmental conditions and genetic background.

### Physiological Stress from Hypoxia Does Not Necessarily Translate to Changes in Recombination

Temperature had a clear effect on recombination across the intervals examined, consistent with previous work showing that both increases and decreases in temperature can alter recombination rates and that these responses often depend on genetic background (Aggarwal et al. 2015, 2019; Belmonte-Tebar et al. 2022; Rybnikov et al. 2023; Novak et al. 2026). Temperature may influence meiotic recombination through changes in processes such as chromatin organization, DNA repair, or the activity of proteins involved in crossover formation (Modliszewski and Copenhaver 2017)(Khaitova et al., 2024; Modliszewski et al., 2018; K. Zhang et al., 2017).

DGRP_57 provides an interesting example of how recombination and physiological responses may not always move together. This background showed one of the strongest increases in recombination under combined low temperature and hypoxia, yet showed relatively small physiological responses to hypoxia in our companion study (Rivera-Rincón et al. 2026). Its respiratory quotient increased by only 0.01 under hypoxia at low temperature, compared with increases of up to 0.12 in most other backgrounds, and CTmax decreased by only 0.13°C under combined stress. This contrast suggests that the physiological response to a stressor does not necessarily predict the magnitude of its effect on recombination.

Hypoxia on its own had a much weaker effect on recombination. This may partly reflect the natural tolerance of *D. melanogaster* to low-oxygen environments during development (Zhou et al. 2008; Burggren et al. 2017; Polan et al. 2020). Larvae and pupae commonly experience reduced oxygen availability in their natural substrates, and adaptations to these conditions may help buffer meiotic processes against moderate oxygen limitation (Zhou et al. 2007, 2008, 2009).

Importantly, the weak recombination response to hypoxia does not mean that hypoxia was physiologically unimportant. In our companion study, hypoxia affected several measures of metabolism, thermal tolerance, body mass, and fertility across the same genetic backgrounds (Rivera-Rincón et al. 2026). Respiratory quotient and CO_2_ production changed under hypoxia, although the magnitude and direction of these responses differed among backgrounds. Thermal tolerance and body mass also responded differently depending on genotype and temperature. These results show that hypoxia imposed measurable physiological effects even when recombination changed relatively little.

Fertility showed a similar disconnect. F_1_ fertility was significantly affected by temperature, oxygen, genetic background, and the interaction between temperature and oxygen, whereas F_2_ fertility was primarily influenced by genetic background (Rivera-Rincón et al. 2026). Thus, reproductive and physiological traits responded strongly to the experimental conditions, while meiotic recombination was comparatively less sensitive to hypoxia alone.

Taken together, these patterns suggest that temperature was the stronger environmental driver of recombination rate plasticity in this experiment. Hypoxia clearly affected other aspects of physiology and reproduction, but its independent effect on recombination was limited. Its influence on recombination became more apparent in particular genetic backgrounds and when combined with low temperature, suggesting that the meiotic response to oxygen limitation depends strongly on environmental and genetic context.

### Genetic Background Shapes the Response to Environmental Stress

The differences among genetic backgrounds were among the clearest patterns in this study. Previous work has shown that genetic variation can influence crossover frequency, interference, and the genomic distribution of recombination (Jeffreys and Neumann 2005; Paigen and Petkov 2010; Stevison et al. 2017; Szymanska-Lejman et al. 2023). Our results extend this pattern by showing that genetic background also influences how recombination responds to environmental stress.

DGRP_57 and DGRP_508 showed the largest increases in recombination under combined hypoxia and low temperature, whereas DGRP_391 showed a more moderate response, DGPR_42 showed relatively small and sometimes oppositely directed changes, and DGRP_491 generally showed little change or decreases. These differences suggest that the same environmental conditions can produce very different recombination responses depending on genotype. Interestingly, the magnitude of recombination plasticity also appeared to be related to baseline recombination rate. DGRP_57 and DGRP_508, which had among the lowest baseline recombination rates, showed some of the largest treatment-associated increases, whereas DGRP_391 and DGRP_491 had relatively high baseline rates and showed comparatively little plasticity (Table 2). This pattern was not universal, however, as DGRP_42 also had a relatively low baseline recombination rate but showed only modest changes, in some cases in the opposite direction. However, in another study, this strain showed significant plasticity to changes in caloric density (Novak et al. 2026). Thus, baseline recombination rate appears to somewhat constrain the magnitude of recombination plasticity, with variation across perturbations. The apparent tendency for backgrounds with lower baseline rates to show larger increases under stress warrants further investigation.

Variation in pathways involved in DNA repair, meiotic regulation, chromatin remodeling, or cellular stress responses could contribute to these differences (Osley et al. 2007; Banerjee and Roy 2021). Previous work has also shown that Drosophila genetic backgrounds differ in their metabolic and cellular responses to environmental stress (Aggarwal et al. 2015). The variation observed here may therefore reflect broader genetic differences in how individual strains respond to challenging environmental conditions.

Genotype-dependent recombination responses are not limited to Drosophila. Studies in *Arabidopsis thaliana* and *Caenorhabditis elegans* have also found that environmental conditions can alter recombination differently among genetic backgrounds (Crombie et al. 2019; Zhao et al. 2024). Together, these studies emphasize that genetic context is an important part of understanding recombination plasticity and may help explain why populations differ in their responses to environmental change.

## Conclusions

This study showed that recombination in *Drosophila melanogaster* responds to environmental conditions, but that the response depends strongly on both the type of stressor and genetic background. Low temperature had the clearest overall effect on recombination, whereas hypoxia alone produced relatively limited changes. When hypoxia and low temperature stresses were combined together, however, some genetic backgrounds showed larger shifts in recombination, particularly within the *cv-v* and *v-f* intervals.

The five genetic backgrounds responded differently in both the magnitude and direction of these changes. DGRP_57 and DGRP_508 showed some of the strongest increases under combined stress, whereas DGRP_491 generally showed decreases or little change. These differences emphasize that recombination responses to environmental stress cannot be predicted from environmental conditions alone.

Although hypoxia had clear physiological and reproductive effects, these changes did not consistently translate into altered recombination. This disconnect suggests that different biological traits may respond to the same environmental stress through different mechanisms and with different levels of sensitivity. We also found that baseline recombination may be a better predictor than organismal stress response for recombination rate plasticity.

Overall, our results show that recombination is shaped by an interaction between genetic and environmental variation. Understanding that variation will be important for predicting how recombination patterns may change as organisms encounter increasingly variable and complex environmental conditions.

## Acknowledgments

We would like to thank Dr. Hood, Dr. Schwartz, Dr. Bernal, and Dr. Stoeckel for their revisions and feedback on the manuscript. We would like to thank members of the Stevison Lab for helpful feedback throughout this project. Research reported in this publication was supported by the National Institute of General Medical Sciences of the National Institutes of Health under Award Number R35GM147501. The content is solely the responsibility of the authors and does not necessarily represent the official views of the National Institutes of Health.

## Statements and Declarations

Declaration of competing interest: The authors declare that they have no known competing financial interests or personal relationships that could have appeared to influence the work reported in this paper.

## Author Contributions

Conceptualization Laurie Stevison and Natalia Rivera-Rincón]; Methodology Natalia Rivera-Rincón, Emma Saurette, Logan Havard; Formal analysis and investigation Laurie Stevison and Natalia Rivera-Rincón; Writing – original draft preparation Laurie Stevison and Natalia Rivera-Rincón; Writing – review and editing Natalia Rivera-Rincón, Emma Saurette, Logan Havard, Laurie Stevison; Funding acquisition Laurie Stevison; Resources Laurie Stevison; Supervision Laurie Stevison and Natalia Rivera-Rincón

## Notes

https://github.com/StevisonLab/Recombination_Hypoxia_LowTemp_in_DMEL

### Competing Interest Statement

The authors have declared no competing interest.

