## Supplement for "Interaction of combined stressors and genetic background on recombination rate plasticity in *Drosophila melanogaster*"

### Supplementary Figures

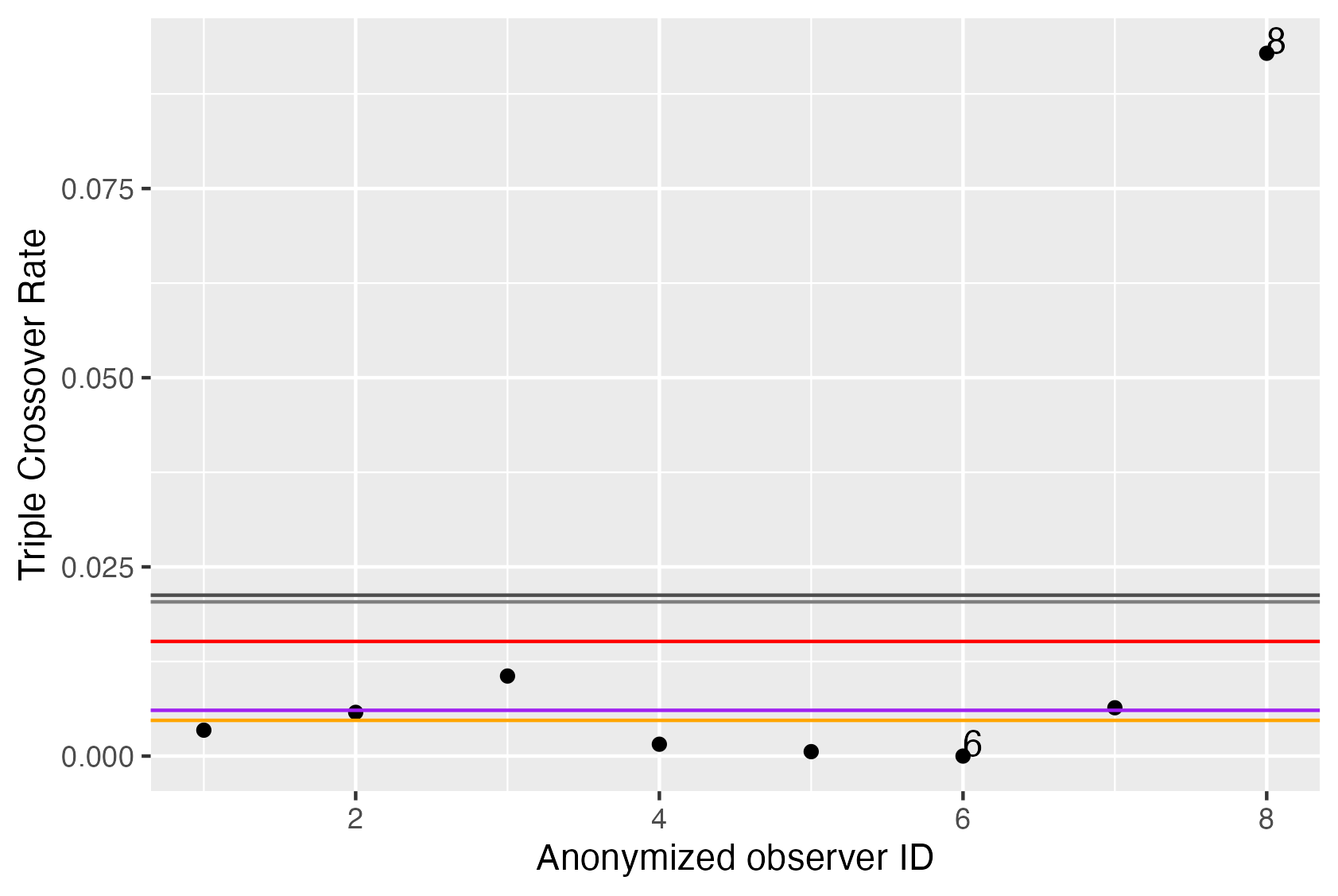

**Figure 1. Observer Bias Analysis.** Here we compare the rate of triple crossovers (TCOs) between 8 total observers in the full dataset. The mean across all observers is shown in red, with the expected rate based on the map lengths from FlyBase shown in Purple. The mean plus the median absolute deviation (MAD) is shown in light grey with the mean plus the interquartile range (IQR) shown in dark grey. Based on observed TCO rates in excess of both the MAD and IQR, all observations from individual 8 were removed from the dataset. Similarly, all observations from individual 6 were removed due to fewer than 50 total male observations. The adjusted mean TCO rate of the remaining 6 observers is shown in orange. This analysis was similar to previous work using the same set of individuals and analysis. It is worth noting that though these results are anonymized here, the same individual had elevated TCO rates across datasets, suggesting this represents a real bias in the set of observations for this individual across studies.

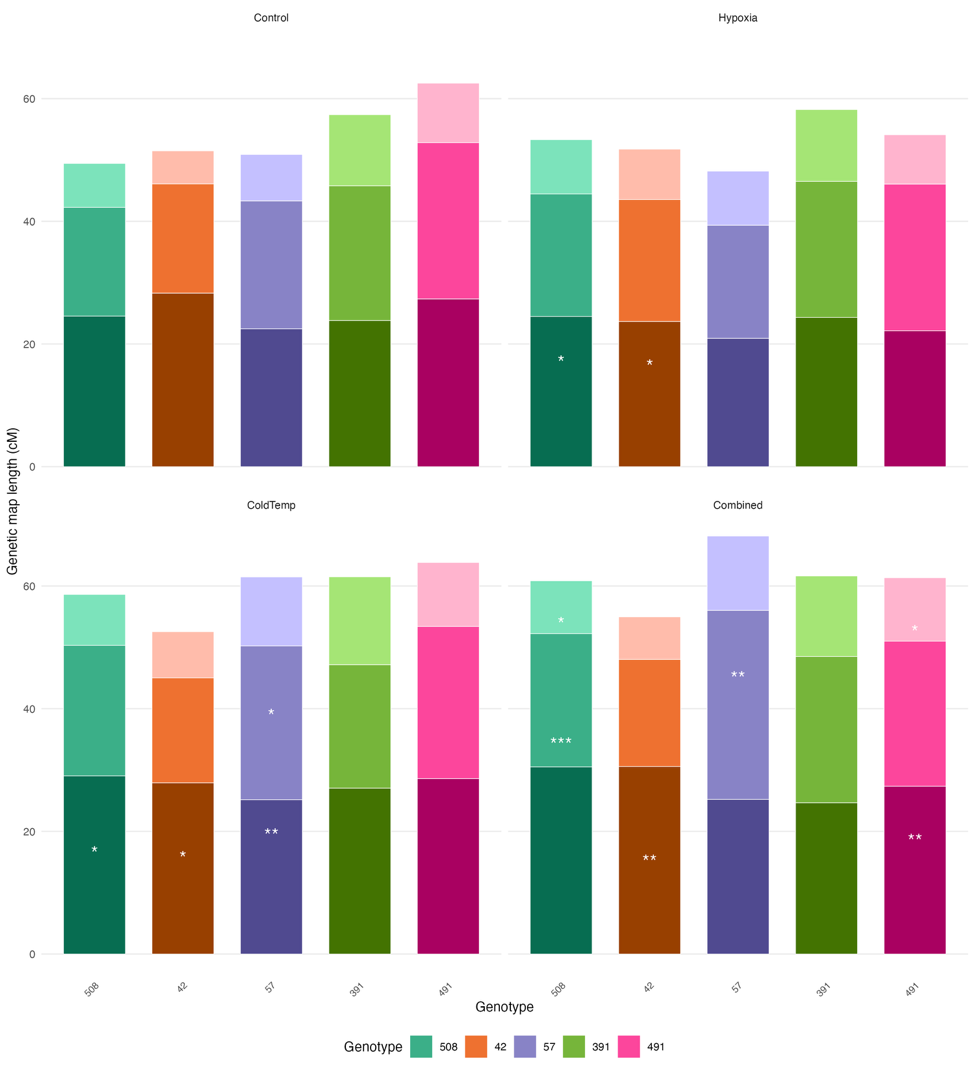

**Figure 2. Recombination results per interval across all five genotypes and four treatments.** Each panel shows a treatment group with genetic map lengths shown as a stacked bar plot. Each bar represents a different genotype. The height of each bar represents the total genetic map length (cM) across the full y-f interval. Line plot showing only total map length can also be found in Supplemental figure 2. Significance of each interval indicated by asterisks. Full posthoc results by time point for each treatment contrast can be found in Supplemental figures 3-7. An expanded plot per time point of this figure is in Supplemental figure 8.

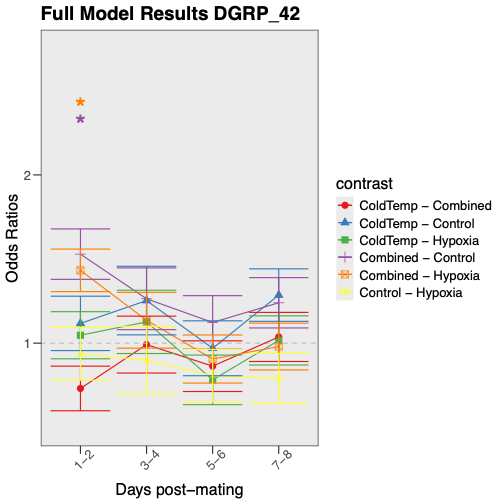

**Figure 3. Recombination Rate Results for DGRP_42.** Based on the significant 3-way interaction term (Table 3), a post hoc analysis was done for treatment, genetic background and time point. The coefficients from the logistic regression model were exponentiated to calculate the odds ratio for each treatment contrast for each genetic background and for each timepoint. Significant contrasts between each of the four treatment groups are indicated by asterisks. Overall, recombination rate in DGRP_42 was significantly higher in the combined treatment relative to both the control and hypoxia treatments in days 1-2 of the experiment.

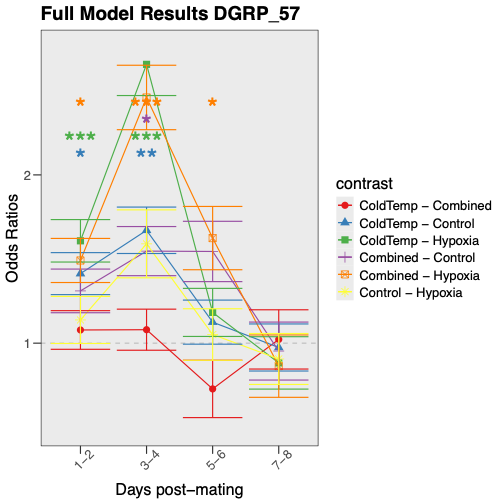

**Figure 4.Recombination Rate Results for DGRP_57.** As in Figure 3 above, this plot is of the posthoc treatment contrasts for the 3-way interaction term. Here, recombination rate in DGRP_57 was significantly lower in hypoxia vs. combined in three of the four timepoints. Additionally, coldtemp was significantly higher than both control and hypoxia in the first four days of the experiment. Finally, combined was significantly higher than control on days 3-4.

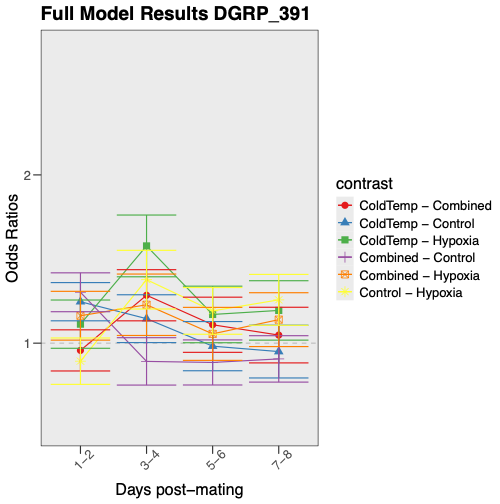

**Figure 5. Recombination Rate Results for DGRP_391.** As in Figure 3 above, this plot is of the posthoc treatment contrasts for the 3-way interaction term. Here, DGRP_391 had no significant differences between any treatment comparisons for any of the timepoints.

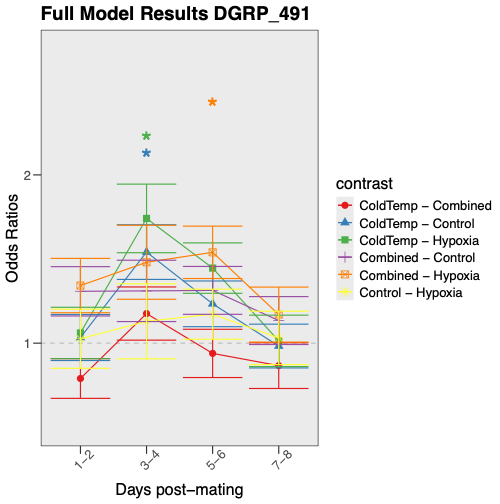

**Figure 6. Recombination Rate Results for DGRP_491.** As in Figure 3 above, this plot is of the posthoc treatment contrasts for the 3-way interaction term. Here, DGRP_491 had significant contrasts between control vs. coldtemp as well as hypoxia vs. coldtemp on days 3-4. Additionally, hypoxia vs. combined was significantly different on days 5-7.

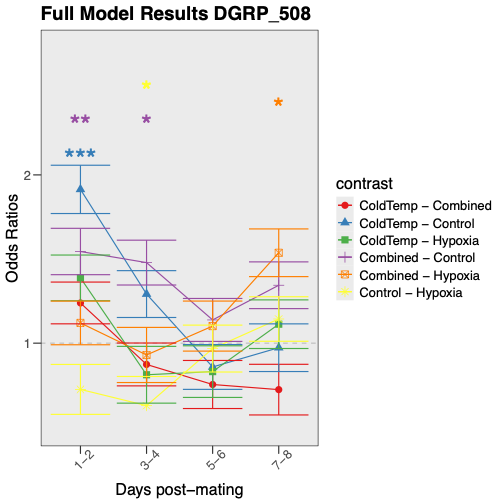

**Figure 7. Recombination Rate Results for DGRP_508.** As in Figure 3 above, this plot is of the posthoc treatment contrasts for the 3-way interaction term. Here, DGRP_508 had significant contrasts between control vs. combined in the first half of the experiment. Additionally control vs. coldtemp was different on days 1-2, control vs. hypoxia was different on days 3-4, and hypoxia vs. combined was different on days 7-8.

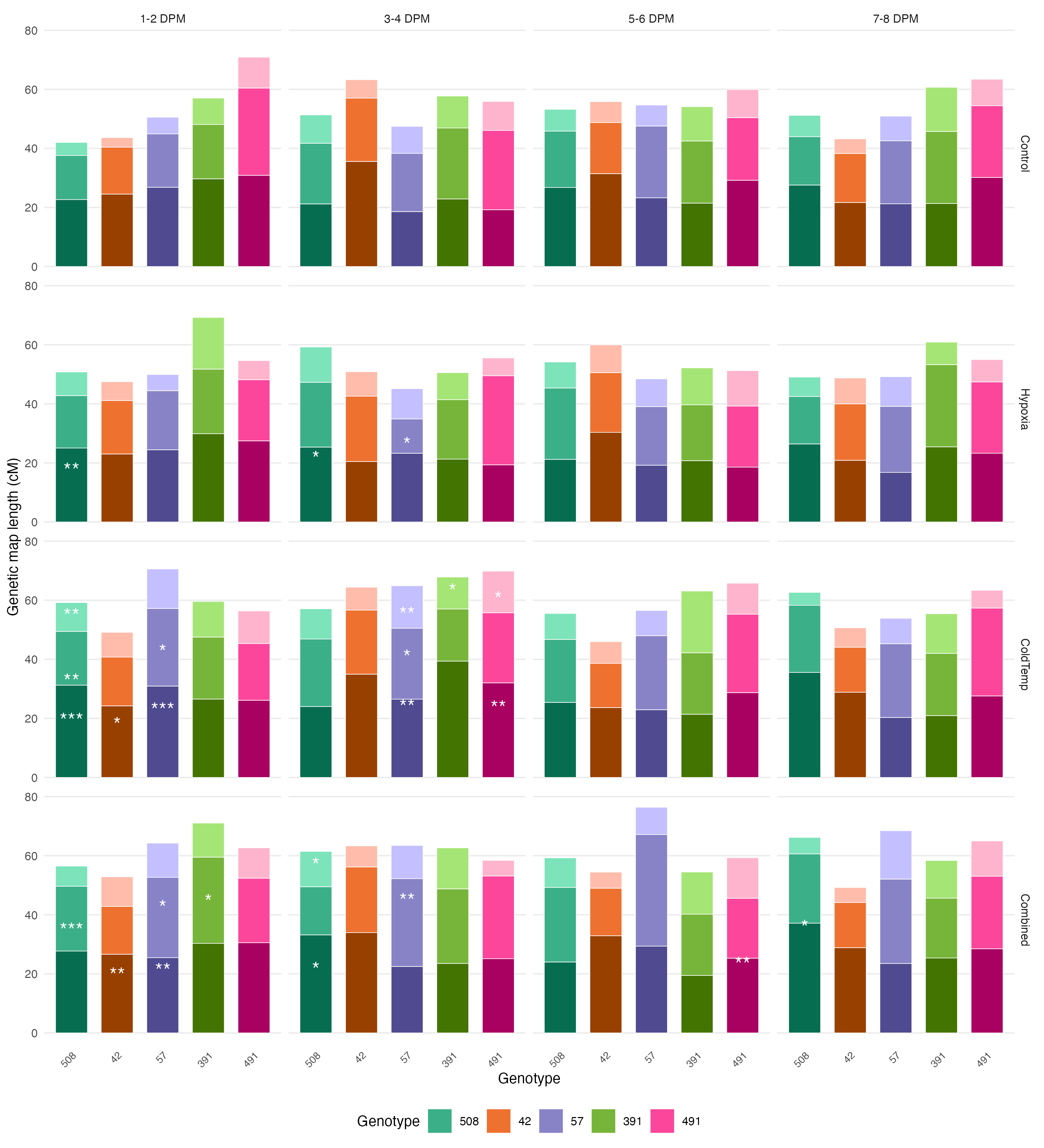

### Supplementary Tables

Table 1. Model Results when treatment split into two parameters. Here, temperature and oxygen were coded as separate model parameters each with two levels. An AIC comparison showed this model was equivalent to a model with a single treatment parameter with four levels, which is what is presented in Table 4 in the main text. To facilitate comparisons across treatment groups and to opt for a simpler model with fewer parameters, the single treatment model was selected.

| Model Parameter | Df | Chi-Square | r_1-2-3_  *y-cv-v-f* |
| --- | --- | --- | --- |
| Oxygen | 1 | 0.85 | 0.36 |
| Temperature | **1** | **39.36** | **3.53e-10** |
| Genetic Background (GB) | **4** | **12.68** | **0.013** |
| Brood | 3 | 5.99 | 0.11 |
| Oxygen:Temperature | 1 | 0.94 | 0.33 |
| Oxygen:GB | **4** | **9.56** | **0.048** |
| Temperature:GB | 4 | 6.41 | 0.17 |
| Oxygen:Brood | 3 | 4.08 | 0.25 |
| Temperature:Brood | **3** | **21.48** | **8.37e-05** |
| GB:Brood | **12** | **57.44** | **6.57e-08** |
| Oxygen:Temperature:GB | 4 | 2.49 | 0.65 |
| Oxygen:Temperature:Brood | 3 | 3.55 | 0.31 |
| Oxygen:GB:Brood | **12** | **21.97** | **0.038** |
| Temperature:GB:Brood | **12** | **30.54** | **0.0023** |
| Oxygen:Temperature:GB:Brood | **12** | **27.27** | **0.0071** |
